# STAT3 signalling enhances tissue expansion during postimplantation mouse development

**DOI:** 10.1101/2024.10.11.617785

**Authors:** Takuya Azami, Bart Theeuwes, Mai-Linh N Ton, William Mansfield, Luke Harland, Masaki Kinoshita, Berthold Gottgens, Jennifer Nichols

**Affiliations:** Cambridge Stem Cell Institute, University of Cambridge, Jeffrey Cheah Biomedical Centre, Puddicombe Way, Cambridge CB2 0AW, UK; Department of Physiology, Development and Neuroscience, University of Cambridge, Tennis Court Road, Cambridge CB2 3EG, UK; Centre for Trophoblast Research, University of Cambridge, UK; MRC Human Genetics Unit, Institute of Genetics and Cancer, University of Edinburgh, Western General Hospital, Crewe Road South, Edinburgh EH4 2XU, UK; Babraham Institute, Babraham Hall House, Cambridge CB22 3AT, UK; School of Biosciences, University of Nottingham, Sutton Bonington Campus, Nottingham, LE12 5RD, UK

**Keywords:** Chimeras, Developmental delay, Pluripotent stem cells, RNA sequencing, STAT3 signalling

## Abstract

STAT3 signalling has been studied extensively in the context of self-renewal and differentiation of mouse embryonic stem cells. Zygotic STAT3 is required for normal postimplantation development. On an outbred genetic background, *Stat3* null embryos consistently lagged behind their littermates, beginning with significant reduction of epiblast cells at implantation. Remarkably, mutants closely resemble non-affected embryos from the previous day at all postimplantation stages examined. We pinpoint this phenotype to loss of the serine-phosphorylated form of STAT3 which predominates in postimplantation embryonic tissues. Bulk RNA-sequencing analysis of isolated mouse epiblasts confirmed *Stat3* null embryos exhibited developmental delay transcriptionally. Single cell RNA sequencing of mid gestation chimaeras containing STAT3 null embryonic stem cells revealed exclusion of mutant cells exclusively from the erythroid lineage. Although Stat3 null embryonic stem cells can differentiate into erythroid and hematopoietic lineages in vitro, they are out-competed when mixed with wild type cells. Combined with the reduced size of STAT3 null epiblasts after implantation, our results implicate a role for STAT3 in cell proliferation affecting temporal control of embryonic progression and rapid differentiation.

For the purpose of Open Access, the author has applied a CC BY public copyright licence to any Author Accepted Manuscript version arising from this submission.

## Introduction

Signal Transducer and Activator of Transcription (STAT) 3 is stimulated by the cytokine Leukaemia Inhibitory Factor (LIF) in murine embryonic stem cells (ESCs) via activation of Janus-associated kinases (JAKs) inducing phosphorylation of STAT3 at tyrosine 705 (pY705) (Ni *et al*, 2004; Zhang *et al*, 2000). STAT3 can also be phosphorylated at serine 727 (pS727) by mitogen-activated protein kinases (MAPK) (Huang *et al*, 2014). Mutation of pY705 ablates ESC self-renewal in standard (serum/LIF) culture conditions, whereas pS727 is apparently not needed in ESCs prior to neuronal differentiation (Huang *et al*., 2014). A requirement for pS727 was reported for mitochondrial STAT3 transcriptional activity in cell lines (Carbognin *et al*, 2016; Gough *et al*, 2013; Peron *et al*, 2021), suggestive of a role in metabolic function during differentiation. Zygotic deletion of exons 20-22 of the *Stat3* gene produces embryos that can implant in the uterus but further development is impaired (Takeda *et al*, 1997). In blastocysts, STAT3 regulates cellular metabolism in mitochondria by promoting oxidative phosphorylation and expression of naïve pluripotency-associated genes (Betto *et al*, 2021), but the role of STAT3 at post-implantation stages is largely unknown.

In this study, we outcrossed *Stat3* heterozygous mice (Takeda *et al*., 1997) to the robust CD1 genetic background and examined postimplantation development. Intriguingly, *Stat3* null embryos could survive beyond gastrulation, but exhibited defects in peri-implantation epiblast expansion and temporal control of embryonic progression and metabolism. Using our recently derived *Stat3* null ESCs (Kraunsoe *et al*, 2023), we compared proliferation dynamics in self-renewing ‘2i’ culture conditions, comprising inhibition of MEK/ERK and GSK3, with wild type ESCs derived from the same genetic background. To scrutinise the tissue-forming capacity of embryonic cells lacking STAT3 during development we generated chimaeras by injecting *Stat3* null ESCs into WT CD1 host blastocysts and analysed their ability to contribute to emerging tissues by means of single cell chimaera sequencing using an updated version of our previously optimised protocol (Guibentif *et al*, 2021; Pijuan-Sala *et al*, 2019). Although able to colonise most tissues and coordinate developmental stage with the host embryo, *Stat3* null cells were specifically excluded from the primitive blood lineage, consistent with a requirement for STAT3 signalling in tissues associated with rapid proliferation.

## Results

### STAT3 is dispensable for organogenesis but required to sustain developmental pace

We recently showed that mouse embryos lacking *Stat3* rapidly lose their inner cell mass (ICM) component following induction of embryonic diapause (Kraunsoe *et al*., 2023). Here, we examined non-diapause embryos generated by inter-crossing *Stat3* heterozygous mice on the CD1 genetic background to determine at which stage developmental deficiencies might arise. We first compared peri-implantation embryonic day (E) 4.5 *Stat3* null versus wild type (WT) or heterozygous (het) embryos using immunofluorescence (IF) for the epiblast marker, NANOG, and primitive endoderm (PrE) marker, GATA6. Significantly fewer NANOG+ cells were observed in *Stat3* null embryos, whereas the numbers of GATA6+ cells did not differ appreciably between genotypes (Fig. 1A,B). A dramatic reduction in the mean number of trophectoderm cells, identified by absence of NANOG and GATA6 and occupation of the outer layer, in mutant embryos compared with WT or hets was observed (Fig. 1B). We attribute this phenotype to decreased FGF signalling resulting from the small null epiblasts, as predicted by previous observations implying requirement for FGF4 in trophectoderm proliferation (Nichols *et al*, 2009; Nichols *et al*, 1998).

**Figure 1.**
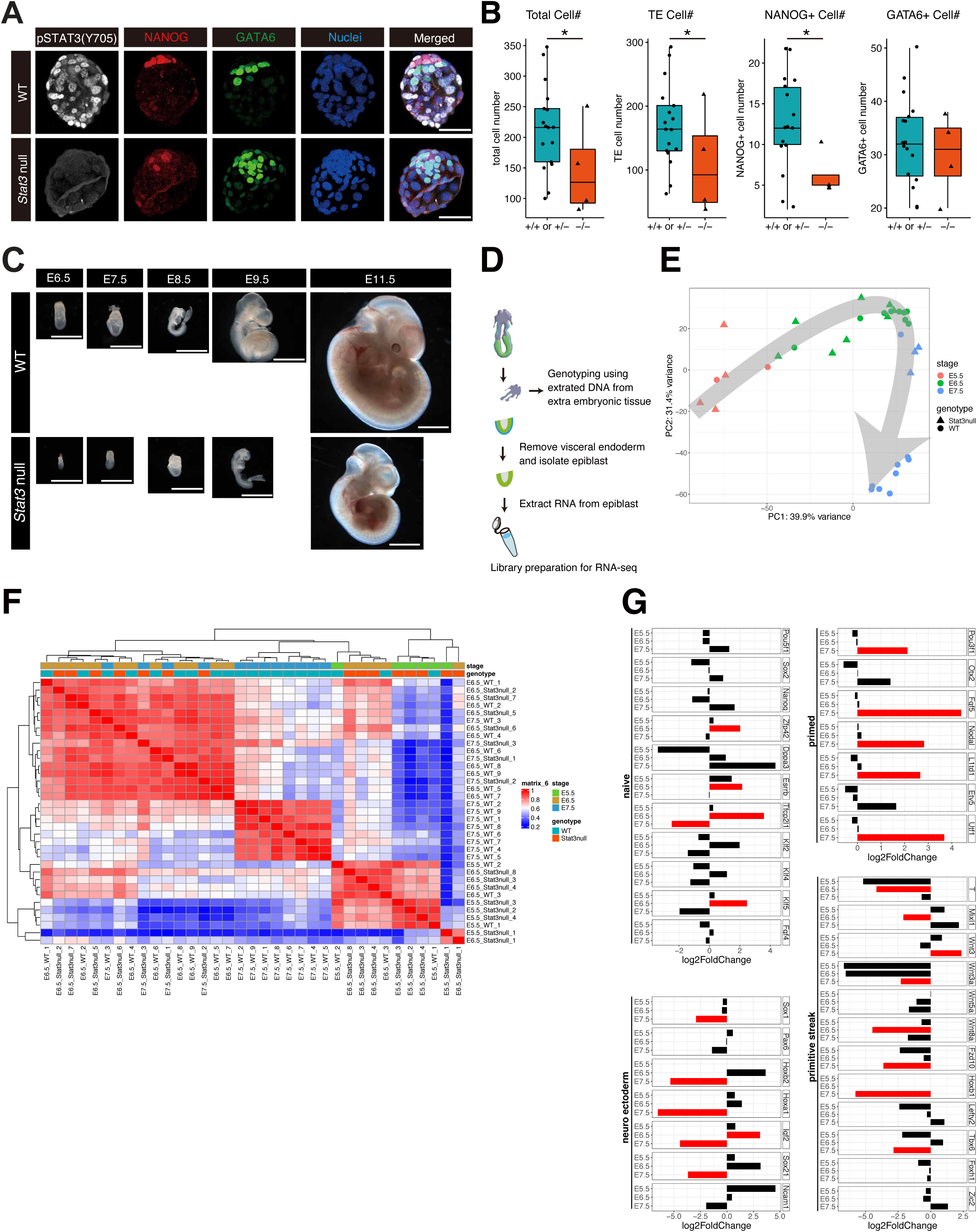
Developmental retardation of *Stat3* null embryos. (A) Immunofluorescence for pSTAT3(Y705), NANOG, and GATA6 in E4.5 WT and *Stat3* null embryos. Scale bar = 50 µm. (B) Quantification of NANOG+ and GATA6+ cells in (A) WT: n = 17, *Stat3* null: n = 4, * *P* < 0.01. (C) Morphology of *Stat3* WT and null embryos from E6.5 to E11.5. Scale bar = 1 mm. (D) Schematic of bulk RNA-sequencing of epiblasts isolated from postimplantation embryos. (E) PCA plot of WT and *Stat3* null epiblast cells from E5.5 to E7.5. (F) Heatmap comparison and unsupervised hierarchical clustering of WT and *Stat3* null epiblast cells from E5.5 to E7.5. (G) Log2 fold change comparison for naïve, primed, neuroectoderm, and primitive streak genes. Red bar indicates significant differences (adjusted p < 0.05).

Surprisingly, on the CD1 genetic background, *Stat3* null embryos could be retrieved as late as E11.5 (Table S1), but consistently exhibited a phenotype more closely resembling the morphology expected for the previous day of development compared with their littermates, with no other obvious morphological defects (Fig. 1C; Fig. S1). This intriguing observation implies that embryos lacking STAT3 are unable to instigate sufficient cell division soon after implantation, when mechanisms for size regulation are normally evoked (Rands, 1986; Snow, 1976). To our knowledge, the sustained period of developmental delay we have observed lacking overt signs of abnormalities in organogenesis has not been reported previously.

### STAT3 null epiblasts follow a normal, but delayed developmental trajectory

Epiblasts were dissected from CD1 *Stat3* het *inter-se* mating at E5.5, E6.5 and E7.5, thus spanning epiblast transition from naïve, via formative (Kinoshita *et al*, 2020) to primed pluripotency (Nichols & Smith, 2009) and the onset of gastrulation. Each embryo was genotyped using extraembryonic tissue (Fig. 1D). From E5.5 to E6.5 the epiblast is basically a homogeneous epithelium poised to respond to asymmetric signals from the extraembryonic tissues. RNA-sequencing (RNAseq) was therefore performed in bulk on denuded epiblasts to compare transcription profiles between *Stat3* WT and null epiblasts. While E5.5 WT and *Stat3* null samples exhibited very similar transcriptional identity, E6.5 null epiblasts were clearly retarded, with an expression pattern intermediate between WT E5.5 and E6.5 (Fig. 1E,F). E7.5 null epiblasts grouped more closely with WT E6.5 than E7.5 epiblasts by principal component analysis (PCA; Fig. 1E) and unsupervised hierarchical clustering (Fig. 1F), suggesting that null epiblasts have not yet advanced from formative to primed pluripotency. *Stat3* null epiblasts exhibited downregulation of naïve and upregulation of primed pluripotency genes at E6.5 (Fig. 1G), concurrent with preparation for primitive streak formation *in vivo* (Snow, 1976). However, mutants failed to induce neuroectoderm significantly and displayed reduced intensity of selected primitive streak genes at E7.5, consistent with the temporal delay of gastrulation (Fig. 1G).

### STAT3 activity is controlled by phosphorylation at Serine 727 in postimplantation embryos

Validated antibodies for each of the two STAT3 phosphorylation forms (pY705 and pS727) were used for IF studies on WT CD1 mouse embryos from implantation until gastrulation. As previously shown (Kraunsoe *et al*., 2023), nuclear phosphorylated-Y705 STAT3 was detected in peri-implantation embryos at E4.5 (Fig. 1A). Apart from the extra-embryonic visceral endoderm, pY705 was barely detectable in WT embryos soon after implantation, but re-emerged in the node region of the primitive streak at mid-late gastrulation (Fig. 2A). The dramatic reduction of pY705 in postimplantation epiblasts correlates with the low level of *Socs3* revealed in the bulk RNAseq (Fig. 2B). In contrast, pS727 was abundant in the epiblast, extra-embryonic ectoderm and extra-embryonic endoderm from E4.5 to E6.5, becoming restricted to visceral endoderm, epiblast and its derivatives at E7.5 (Fig. 2C). These stages coincide with the requirement for elevated mitochondrial function in the embryo to enable rapid cell division for tissue expansion during gastrulation (Lima *et al*, 2021). To identify potential signalling pathways activated by pS727, we applied candidate inhibitors to WT ESCs to test the phosphorylation reaction *in vitro*. MEK inhibition by PD0325901 (PD03) to block FGF signalling in serum-free medium did not significantly impair phosphorylation of Y705 or S727 (Fig. 2D,E), whereas S727 phosphorylation was previously shown to depend upon ERK signalling in serum-containing medium (Huang *et al*., 2014). Interestingly, JAK inhibition ablated Y705 phosphorylation, but only partially dampened pS727, indicating that S727 can be phosphorylated by factors distinct or downstream from JAK signalling or LIF-gp130 receptors (Fig. 2D,E).

**Figure 2.**
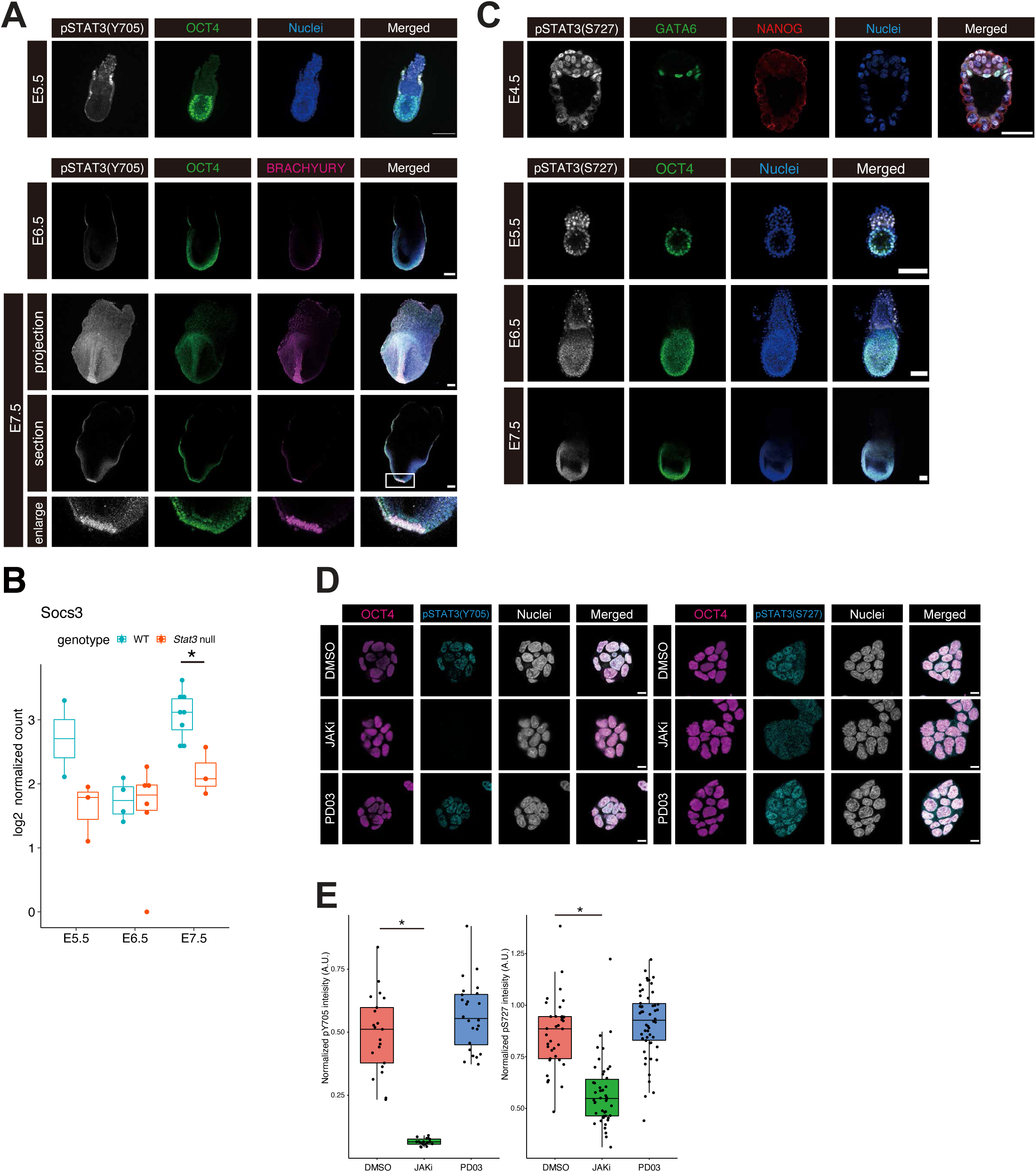
The predominant active form of STAT3 transitions from pY705 to pS727 in WT embryonic lineages after implantation. (A) Immunofluorescence for pSTAT3(Y705) and OCT4 in E5.5 embryos and pSTAT3(Y705), OCT4, and BRACHYURY in E6.5 and E7.5 embryos. Scale bar = 100 µm. (B) log2 normalized expression levels of *Socs3* in WT and *Stat3* null epiblast cells from E5.5 to E7.5. \**P* < 0.01. (C) Immunofluorescence for pSTAT3(S727), GATA6, and NANOG in E4.5 embryos and pSTAT3(S727) and OCT4 in E5.5, E6.5, and E7.5 embryos. Scale bar = 25 µm for E4.5 and 100 µm for E5.5, E6.5, and E7.5 embryos. (D) Immunofluorescence for pY705 and pS727 STAT3 in mouse ESCs. ESCs were cultured in N2B27+2i/L for 24 h, then medium exchanged to N2B27 with DMSO, JAK inhibitor, or PD03. Scale bar = 10 µm. (E) Quantification of pY705 and pS727 STAT3 intensity from immunofluorescence. * *P* < 0.01. A.U., arbitrary unit.

### Self-renewing stem cell lines can be derived from STAT3 null postimplantation epiblasts

Despite their reduced size compared with WT littermates, epiblasts from *Stat3* null embryos at E6.5 and E7.5 could be captured and propagated as epiblast stem cell (EpiSC) lines (Brons *et al*, 2007; Tesar *et al*, 2007) with high efficiency (Table S2). Their morphology in culture appeared indistinguishable from that of WT (Fig. 3A) and growth kinetics were similar (Fig. 3B). Transcriptional profiles of *Stat3* WT and null EpiSCs exhibited close clustering by means of bulk RNAseq (Fig. 3C), implicating a STAT3-independent *in vitro* adaptation to self-renewal during acquisition of the mid-gastrulation anterior streak identity reported for EpiSCs using the original culture conditions, regardless of stage of origin (Kojima *et al*, 2014). Differential expression analysis identified 39 upregulated and 99 downregulated genes in *Stat3* null EpiSCs related to WT (Fig. 3D,E). Among the significantly downregulated genes in Stat3 null EpiSCs, the gene ontology (GO) terms for cell migration and cell adhesion were substantially enriched (Fig. 3F). Consistent with the small transcriptional changes in Stat3 null EpiSCs compared to WT, neither pY705 nor pS727 STAT3 was detected in WT EpiSCs (Fig. 3G,H), which was surprising considering the abundant pS727 STAT3 detected in WT postimplantation epiblasts (Fig. 1E). This suggests that, although the FGF-ERK pathway phosphorylates S727 STAT3 in ESCs in serum/LIF medium (Huang *et al*., 2014), pS727 is not induced significantly by FGF in standard, serum-free EpiSC culture. LIF supplementation is not required for EpiSC culture (Brons *et al*., 2007; Tesar *et al*., 2007); however, pY705 and pS727 were both induced by administering LIF to WT EpiSCs (Fig. 3G,H). We conclude that STAT3 signalling is dispensable for maintenance of primed pluripotent cells *in vitro,* which is most likely attributable to relatively slow cell doubling times characteristic of EpiSCs in culture compared with pre-primitive streak epiblast cells *in vivo* (Snow, 1976).

**Figure 3.**
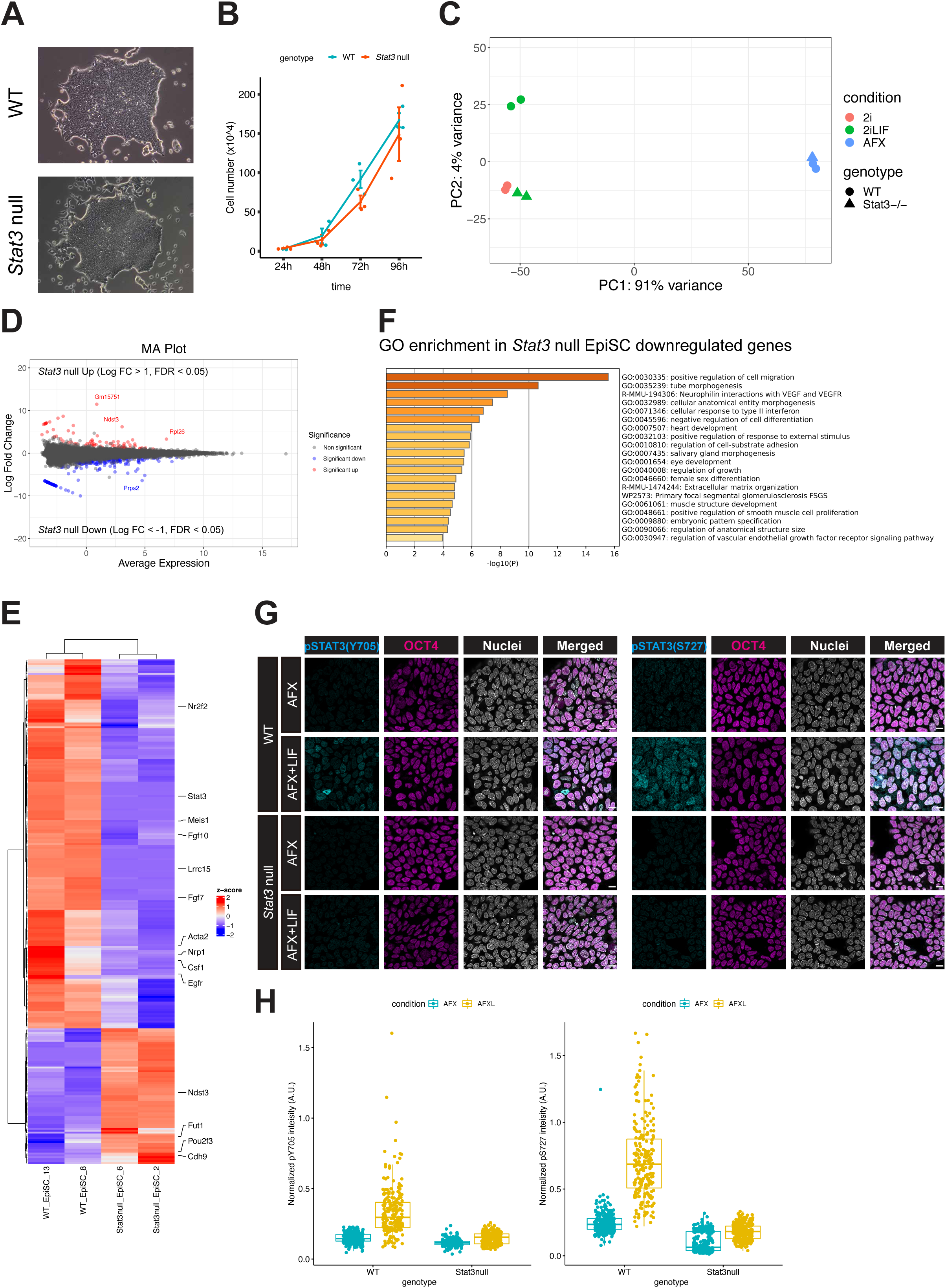
Derivation and analysis of *Stat3* WT and null EpiSC. (A) Representative bright field images of WT and *Stat3* null EpiSCs, derived from epiblast of E6.5 and E7.5 embryos (Table S2). (B) Proliferation of WT and *Stat3* null EpiSCs. (C) PCA plot of WT and *Stat3* null ESCs data obtained from (Betto *et al*., 2021) and EpiSCs from bulk RNA-seq. (D) MA plot of differentially expressed genes (absolute log2 fold change > 1, FDR < 0.05) in *Stat3* null EpiSC related to WT. (E) Heatmap for differentially expressed genes in *Stat3* null EpiSCs compared to WT. (F) Gene ontology (GO) enrichment analysis for the downregulated genes in *Stat3* null EpiSCs identified in (D). Metascape (https://metascape.org/gp/index.html#/main/step1) was used for the GO analysis. (G) Immunofluorescence for pSTAT3(Y705 and S727) and OCT4 in WT EpiSC cultured in N2B27+AFX with or without LIF. Scale bar = 10 µm. (H) Quantification of pSTAT3 intensity from immunofluorescence in (G). * P < 0.01. A.U., arbitrary unit.

### *Stat3* null ESCs are specifically excluded from the erythroid lineage in mid-gestation chimeric embryos

*Stat3* null ESCs were previously shown to contribute to all embryonic lineages upon injection into the blastocyst (Martello *et al*, 2013); however, single cell resolution analyses were not performed to address cell type specific contributions at the transcriptional level. Importantly, in that study, the *Stat3* null ESCs were derived on an MF1 outbred background, whereas the host blastocysts were inbred C57BL/6. This disparity in genetic backgrounds creates a competitive environment, which in this combination favours the donor cells. This is one motivating factor for the traditional choice of C57BL/6 embryos as hosts for efficient generation of genetically modified mouse lines. Notably, differential signal transduction following activation of gp130/LIFR receptors between mouse strains has been reported (Ohtsuka & Niwa, 2015). To avoid discrepancy between donor and host genetic background environments we therefore injected labelled CD1 *Stat3* null ESCs into WT host blastocysts from the same breeding stock to eliminate the risk of encountering confounding inter-strain cell competition effects. We performed single cell chimera sequencing (scChimera-seq) (Guibentif *et al*., 2021; Pijuan-Sala *et al*., 2019) to search for specific differences between *Stat3* null and WT cells within the same embryo and thereby identify tissues that may be compromised by absence of STAT3.

We demonstrated high contribution of *Stat3* null ESC derivatives to E9.5 chimeric embryos (Fig. S2), thereby validating our experimental approach before embarking upon scChimera-seq. *Stat3* null ESCs were injected into CD1 blastocysts, transferred to pseudopregnant recipient mice and dissected at E7.5, E8.5, and E9.5. Embryos exhibiting 30-40% contribution of donor cells (Fig. S3) were selected for analysis, based upon previous optimisation of scChimera-seq (Fig. 4A) (Guibentif *et al*., 2021; Pijuan-Sala *et al*., 2019). They were dissociated and immediately sorted by flow cytometry into WT versus tdTomato-positive *Stat3* null cells. Cell type annotation of scRNA-seq was performed via label transfer from the previously developed extended atlas of mouse gastrulation and early organogenesis (Fig. 4B) (Imaz-Rosshandler *et al*, 2024; Pijuan-Sala *et al*., 2019). *Stat3* null cells exhibited normal overall contribution to chimeras at the single-cell transcriptional level until E8.5. One notable feature of *Stat3* null embryos, revealed by immunofluorescence for the visceral endoderm markers CER1 and FOXA2, is the reduced intensity and abnormal distribution of the cells comprising the anterior visceral endoderm (AVE), an important signalling centre for instructing formation of anterior structures by the underlying epiblast (Fig. S4). In the context of conventional chimeras, the AVE is derived from the host embryo. Support from the WT extra-embryonic tissues provided to the *Stat3* null cells within chimeras might explain why injected cells can apparently keep pace with those of the host embryo in the majority of tissues. However, *Stat3* null cells appeared to be underrepresented in the erythroid lineage (Fig. 4B,C; Fig S5). Pseudotime analysis was performed on erythroid differentiation trajectories in WT and *Stat3* null cells, confirming a significant reduction in contribution to this lineage by *Stat3* null cells in the transition from E8.5 to E9.5 (Fig. 4D). Differential gene expression analysis along the erythroid pseudotime revealed dysregulation of canonical *Stat3* target genes, such as *Cish* and *Pim1*, as well as failure to upregulate *Stat5a* expression (Fig. 4E,F; Fig S5A). Since *Acss1* (acetyl-CoA synthases) and *Lyrm7* (complex III assembly factor) have been shown to function in mitochondrial energy synthesis (Maio *et al*, 2017; Milger *et al*, 2006), their reduced expression in *Stat3* null erythroid cells may restrict their contribution to this normally rapidly proliferating lineage (Fig. 4E,F).

**Figure 4.**
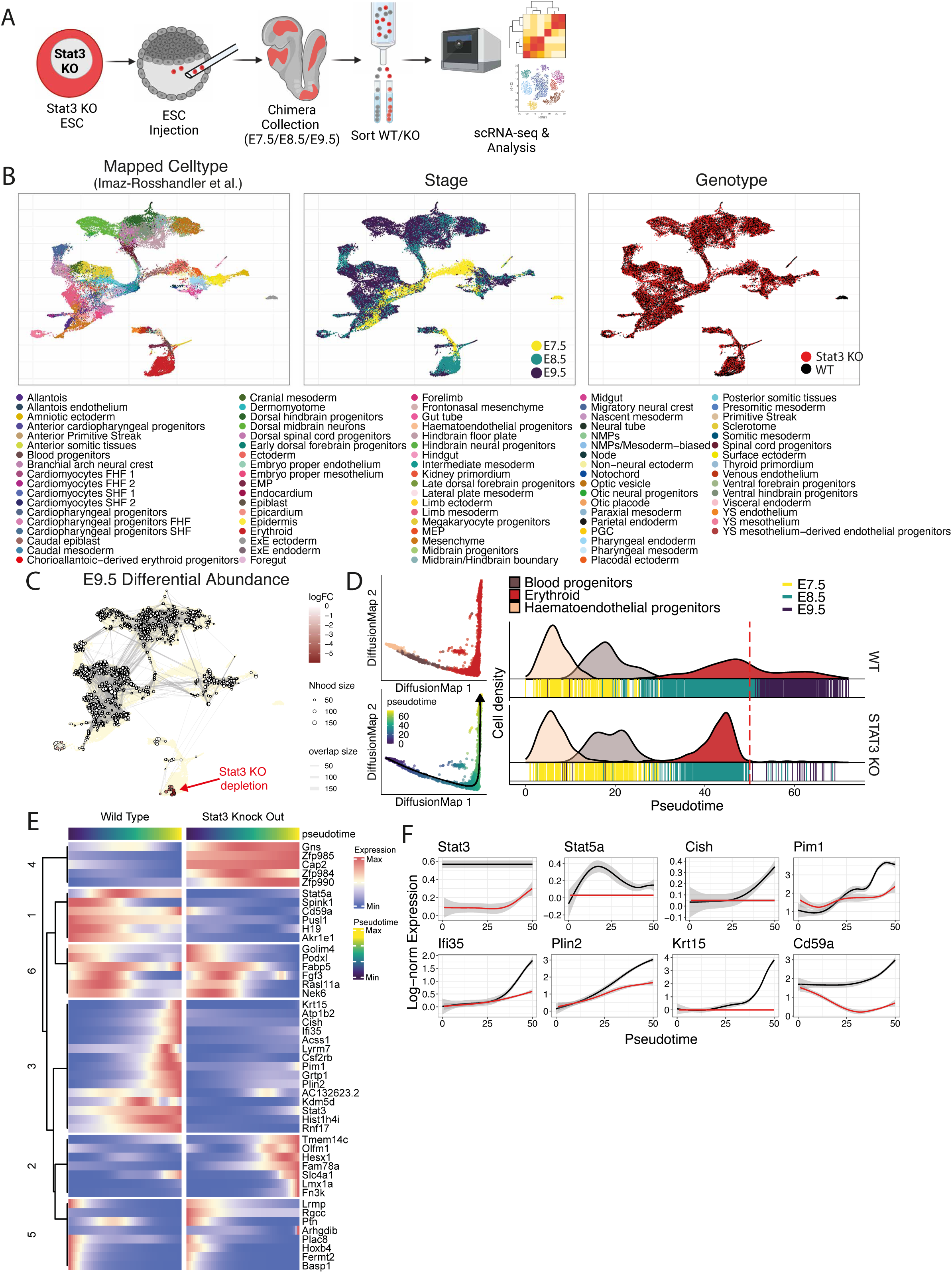
Stat3 null cells are depleted during erythroid lineage differentiation *in vivo*. (A) Experimental design for generation of Stat3 null/WT chimaera-seq experiments. Created with BioRender.com. (B) UMAP representing transcriptional landscape of single-cells from chimaera-seq experiment coloured by label transferred cell type (left), Embryonic stage (middle), and genotype (right). (C) Differential abundance between equivalent *Stat3* KO and WT cells of E9.5 chimera. Negative logFC indicates a loss of Stat3 KO cells in a specific cellular neighbourhood. (D) DiffusionMap of Haematoendothelial progenitor to Erythroid differentiation (left) coloured by label transferred cell type (top) and pseudotime (bottom). Cell density along pseudotime for WT and *Stat3* KO (right). Red dotted line indicates pseudotime cutoff used for downstream analysis. (E) TradeSeq inferred differential genes between WT and *Stat3* KO along Haematoendothelial progenitor to Erythroid differentiation. (F) Log-normalised expression of a subset of genes along pseudotime, for *Stat3* KO (red) and WT (black).

### Differentiation of Stat3 null ESCs to primitive erythroid cells is impeded when co-cultured with WT cells

To investigate further the potential of *Stat3* null cells to differentiate into hematopoietic and primitive erythroid lineages, we applied a recently developed *in vitro* differentiation protocol (Harland *et al*, 2021), which induces yolk sac-like hematopoietic and endothelial progenitors by supplementing cytokines to embryoid bodies (EBs) (Fig. 5A). When cultured separately, proliferation of *Stat3* null ESCs was equivalent to that of WTs over the differentiation time course, reflecting the relatively prolonged cell cycle duration at this juncture (Fig. 5B). After 4 and 5 days of culture, EBs were dissociated and stained for Flk1 and Pdgfra and the Flk1^hi^/Pdgfra^−^ hematovascular mesoderm population quantified (Fig. 5C). Flow cytometry analysis showed that *Stat3* null ECSs differentiated indistinguishably from WT cells to the hematovascular mesoderm population at day 4 and 5. Further differentiation of EBs to hematopoietic progenitors and erythroid lineages was quantified by CD41^+^/c- Kit^+^ and Ter119^+^/CD71^hi^ respectively. *Stat3* null ESCs differentiated to both primitive erythroid and hematopoietic progenitors with equal efficiency compared to WT, suggesting that the development of these lineages by *Stat3* null cells is not inherently impaired (Fig. 5D; Fig. S6A). This is consistent with our observation that *Stat3* null embryos dissected at E11.5 were not anaemic and contained erythroid cells, although they were retarded by one day compared with their littermates (Fig. 1C; Fig. S1). Following our observation of out-competition of *Stat3* null cells from the erythroid lineage *in vivo* by the WT host embryo, we mixed *Stat3* null ESCs with WTs and performed hematopoietic lineage differentiation on chimeric EBs (Fig. 5E; Fig. S6). The proportions of *Stat3* null to WT cells in mixed EBs maintained initial ratios of 1:1 and 1:4 respectively, but after day 2 *Stat3* null cells became excluded as the EBs embarked upon mesoderm and hematopoietic differentiation (Fig. 5F). Despite their reduced contribution to mixed EBs at day 5, *Stat3* null cells were subsequently able to differentiate into Flk1^hi^/Pdgfra– hematovascular mesoderm, but specifically unable to differentiate into Ter119^+^/CD71^hi^ primitive erythroid cells (Fig. 5G-I; Fig. S7). The failure of *Stat3* null cells to form either of the hematopoietic lineages when competing with the surrounding WT cells is consistent with our discovery of exclusion of *Stat3* null cells from the expanding erythroid lineage in chimeric embryos (Fig. 4).

**Figure 5.**
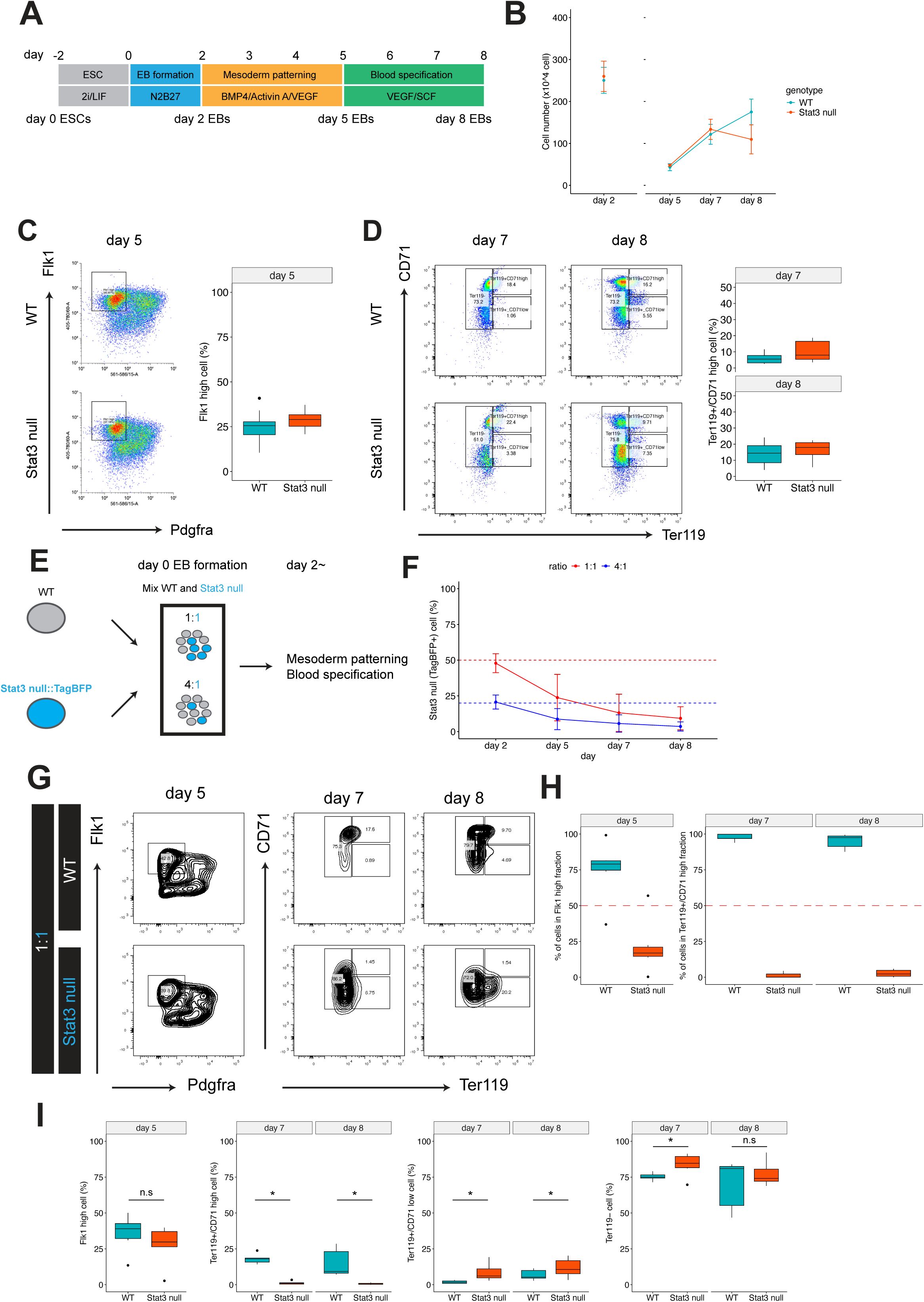
Stat3 null cells can differentiate into hematovascular mesoderm and blood progenitors. Schematics for the hematovascular mesoderm and erythroid differentiation from mouse ESCs. (A) Cell number of embryoid bodies (EBs) differentiated from WT and *Stat3* null ESCs. (C) Representative flow cytometry analysis of Flk1/Pdgfra expression in day 5 WT and *Stat3* null EBs. Hematovascular mesoderm (Flk1^hi^/Pdgfra^−^) are gated in the plot. n = 3 (WT) and n = 3 (*Stat3* null), two independent differentiations. (D) Representative flow cytometry analysis of Ter119/CD71 expression in day 7 and 8 WT and *Stat3* null EBs. Primitive erythroid (Ter119^+^/CD71^hi^) are gated in the plot. n = 8 (WT) and n = 8 (Stat3 null) independent differentiation. (E) Schematics for chimeric EBs differentiation. WT and TagBFP expressing *Stat3* null ESCs were mixed at 1:1 or 4:1 to make EBs. (F) Percentages of *Stat3* null cells in 1:1 and 4:1 mixed EBs over the differentiation. Three WT lines and one TagBFP expressing *Stat3* null ESCs line were used and two independent differentiation experiments were performed and analysed. (G) Representative flow cytometry analysis of Flk1/Pdgfra expression in day 5 and Ter119/CD71 expression in day 7 and day 8 in WT and *Stat3* null cells in chimeric EBs, mixing WT and *Stat3* null ESCs at 1:1. (H) Percentages of WT and *Stat3* null cells in the Flk1^hi^ fraction at day 5, and in the Ter119^+^/CD71^hi^ fraction at day7 and day 8. (I) Percentages of Flk1^hi^ cells at day 5, and Ter119^+^/CD71^hi^, Ter119^+^/CD71^low^, and Ter119^−^ cells at day 7 and day 8 in WT and *Stat3* null cells in chimeric EBs, gated in (G). n.s, not significant. * *p* < 0.05.

## Discussion

*Stat3* null peri-implantation mouse embryos on the CD1 genetic background exhibited a significant reduction in the number of epiblast cells compared with WT counterparts, whereas the number of PrE cells was consistent between both genotypes (Fig. 1). This may be explained by the precocious expression of PrE markers such as *Pdgfra*, *Sox17* and *Gata4* previously observed in *Stat3* null inner cell masses at the mid blastocyst stage (Betto *et al*., 2021) that destabilises the lineage balancing mechanism normally operative within the developing blastocyst (Saiz *et al*, 2016). Previous observations of PrE persistence during diapause when LIFR or gp130 are deleted (Nichols *et al*, 2001) imply independence of this branch of STAT3 signalling for PrE maintenance, whereas deletion of *Stat3* itself resulted in loss of the whole inner cell mass (Kraunsoe *et al*., 2023). Somewhat surprisingly, without the interruption of diapause, *Stat3* null embryos can implant in the uterus and progress through development, but they appear to lack the regulatory mechanisms that allow compensatory proliferation at the onset of gastrulation that would normally enable size regulation in preparation for organogenesis (Rands, 1986; Snow, 1976). The potential consequences of such a defect could be fatal if the number of epiblast cells fails to reach the required threshold, previously shown to be 4 (Morris *et al*, 2012). This phenomenon may explain the relatively high incidence of empty decidua resulting from het inter-crosses and sub-Mendelian proportion of *Stat3* null post-gastrulation embryos recovered (Table S1).

We observed a consistent developmental delay in zygotically deleted *Stat3* mutant post-implantation embryos of approximately one day, reminiscent of the reported role for STAT3 signalling in size regulation in later gestation and post-natal growth (Gutierrez, 2020; Hall *et al*, 2017; Shen *et al*, 2004). *Stat3* null embryos were otherwise apparently normal, suggesting that transition of epiblast from naïve to primed pluripotency (Nichols & Smith, 2009) and subsequent patterning of the fetus can progress, independent of STAT3 signalling (Fig. 1). Consistent with this, propagation of EpiSCs derived from postimplantation epiblast does not require LIF supplementation (Brons *et al*., 2007; Tesar *et al*., 2007), but forced activation of STAT3 or administration of LIF can redirect EpiSCs to an ESC-like state (Bao *et al*, 2009; Yang *et al*, 2010).

Our data reveal that STAT3 pY705 protein is negligible in the WT embryo between implantation and organogenesis, whereas persistent phosphorylation during this period was observed for S727 (Fig. 2), consistent with a previous report implicating requirement for pS727 in exit from pluripotency *in vitro* (Huang *et al*., 2014). Since pS727 is considered to be the mitochondrially-localised form of STAT3 (Carbognin *et al*., 2016; Yang & Rincon, 2016), the reduced growth we observed in *Stat3* null embryos may be a consequence of diminution of mitochondrial function. Bulk RNAseq also confirmed developmental retardation of approximately one day for *Stat3* null epiblasts compared to WT counterparts, but following the same trajectory (Fig. 1). *Stat3* null ESCs injected into WT blastocysts contributed to all tissues until mid-gestation, when exclusion from the erythroid lineage was observed (Fig. 4). This is the first tissue specified at this stage that requires rapid expansion in order to nurture growth and survival of the embryo (Baron *et al*, 2012). In light of the depleted expression of AVE markers CER1 and FOXA2 in extra-embryonic endoderm exhibited by intact *Stat3* null egg cylinders (Fig. S4), we propose that, in the context of a chimera, most *Stat3* null embryonic lineages are at least partially supported by the overlying WT extra-embryonic tissue. Internalization and processing of maternal nutrients are crucial functions of the VE, playing a vital role in facilitating gastrulation (Perea-Gomez *et al*, 2001).

The *in vivo* chimera data we present in Fig. 4 suggest that *Stat3* null cells may be out-competed by WTs specifically when required to differentiate in rapidly proliferating lineages. Our *in vitro* EB differentiation study supports this hypothesis, since *Stat3* null cells were excluded from the hematovascular mesoderm lineage and further erythroid differentiation when co-cultured with WT cells, whereas *Stat3* null cells were able to differentiate along these lineages normally when cultured alone (Fig. 5). Since STAT3 is required for the primitive erythropoiesis via the erythropoietin (EPO) in mouse and zebrafish (Chen *et al*, 2020; Greenfest-Allen *et al*, 2013; Sobah *et al*, 2024), *Stat3* null cells may be expected to exhibit severe defects when differentiated into primitive erythroid cells using EPO. Given that *Stat5a* expression was significantly decreased in *Stat3* null cells in chimeric embryos, competition between WT and *Stat3* null cells for EPO signalling would likely exist, and STAT3 might regulate primitive erythropoiesis by fine-tuning the EPO-STAT5 signalling pathway (Greenfest-Allen *et al*., 2013). The critical aspects of temporal control of embryonic development regulated by the distinct S727 phospho-variant of STAT3 revealed here will inform future studies on erythroid differentiation in the absence of STAT3 and development of protocols to investigate the interaction of embryonic and extra-embryonic tissues during gastrulation regulated by STAT3 signalling.

Mutations in STAT3 signalling are implicated as the cause of various defects in humans, affecting a range of organs and processes. Non-lethal effects include short stature, autoimmunity, and erythropoiesis defects (Mauracher *et al*, 2020), diagnosed at birth and persisting to adulthood (Fabre *et al*, 2019; Kostopoulou *et al*, 2018; Toni *et al*, 2024). Many are the result of point mutations, often resulting in gain of function effects, but the likely consequences of these include disruption to STAT3 function (Gutierrez, 2020). We propose that these human post-natal growth defects may begin during early postimplantation development, resulting from the challenges imposed upon the epiblast to undergo rapid proliferation in preparation for gastrulation. Our discovery that mouse embryos lacking STAT3 can exhibit prolonged developmental delay beyond organogenesis support this hypothesis and provide a novel system with which to dissect mechanisms associated with regulation of proportional organ expansion and overall size regulation.

## Materials and Methods

### Mice, husbandry and embryos

Experiments were performed in accordance with EU guidelines for the care and use of laboratory animals and under the authority of appropriate UK governmental legislation. Use of animals in this project was approved by the Animal Welfare and Ethical Review Body for the University of Cambridge, covered by relevant Home Office licences. Mice were maintained on a lighting regime of 12:12 hours light:dark with food and water supplied *ad libitum*. STAT3 mice heterozygous for replacement of exons 20-22 with Neomycin resistance (Takeda *et al*., 1997) were backcrossed extensively to CD1 mice. Embryos were generated from *Stat3*^+/-^ *inter se* natural mating. Detection of a copulation plug in the morning after mating indicated embryonic day (E) 0.5. Embryos were isolated in M2 medium (Sigma).

### Genotyping

Mice were genotyped by PCR using ear biopsies collected within 4 weeks of birth and genomic DNA was extracted using Extract-N-Amp tissue prep kit (Sigma-Aldrich). Embryos were genotyped using either immune-reactivity to antibody raised against STAT3 pY705 or pS727 in the case of those imaged for confocal analysis, or PCR analysis of trophectoderm lysate for ESC derivation or surplus extraembryonic tissue for postimplantation embryos. Amplification was carried out on around 5 µL of lysate for 35 cycles (following 95°C hot start for 10 minutes) of 94°C, 15 seconds; 60°C, 12 seconds; 72°C, 60 seconds, with a final extension at 72°C for 10 minutes. Reaction products were resolved by agarose gel electrophoresis. Primers used for genotyping PCR are listed in Table.S*3*.

### Histology

Post-gastrulation embryos were fixed in 4% PFA overnight, and serially dehydrated in increasing concentrations of ethanol, rinsed twice in 100% ethanol and embedded in paraffin. Microtome sections of 8 µm thickness were examined histologically via haematoxylin and eosin (H&E) staining.

### Derivation and culture of ESC and EpiSC

Morulae were collected from het females 2.5 days after mating by het males and used for ESC derivation as described previously (Ying *et al*, 2008) by culture to the blastocyst stage in KSOM supplemented with 2i, consisting of 1 µM PD0325901 and 3 µM CHIR99021, transfer of ICMs isolated by immunosurgery to 48-well plates containing 2i in N2B27 medium, one per well. WT and *Stat3* null ESCs were expanded and maintained in N2B27 supplemented with 2i or 2i/LIF on gelatin-coated plates at 37℃ in 7% CO_2_ and passaged by enzymatic disaggregation every 2-3 days. To examine phosphorylation of STAT3 in ESCs, WT cells were cultured in N2B27 with 2i/LIF for 24 h then N2B27 supplemented with DMSO, 1 µM PD0325901, or 0.5 µM JAK inhibitor I (JAKi, Calbiochem) for 2 h. For EpiSC derivation, epiblasts were manually isolated from E6.5 or E7.5 embryos obtained from *Stat3* het inter-cross by removal of extraembryonic ectoderm and visceral endoderm by means of flame-pulled Pasteur pipettes of appropriate diameter. Genotyping was performed using genomic DNA extracted from extraembryonic tissue as described above. Isolated epiblasts were plated on fibronectin-coated plates in N2B27 medium supplemented with AFX, consisting of 12 ng/mL FGF2, 20 ng/mL Activin A and 10 µM XAV929. EpiSCs were maintained in N2B27 supplemented with AFX in fibronectin coated-plates at 37°C in 5% O_2_ and passaged every 3-4 days. To examine phosphorylation of STAT3 in EpiSCs, WT and *Stat3* null EpiSCs were cultured with 10 ng/mL LIF in N2B27 +AFX medium for 1 h.

### Transfection

To generate stable tdTomato and TagBFP-expressing *Stat3* null ESC lines, cells were plated in 24- well plates the day before transfection. 2 µg of plasmid DNA and 2 µL of Lipofectamine2000 (Thermo Fischer Scientific #11668019) were incubated in 25 µL Opti-MEM (gibco) for 5 min, mixed and incubated for 20 min at room temperature. The mixture was added to cells and cultured for 3 hr, then exchanged for fresh medium. After 24 h, cells were passaged to gelatin-coated 6-well plates and puromycin (1 µg/mL) selection performed. Expanded colonies were picked manually and cell lines uniformly expressing tdTomato and TagBFP selected by flow cytometry.

### Blood progenitor differentiation

WT and Stat3 null ESCs were differentiated into mesoderm and blood progenitors as previously described (Harland *et al*., 2021), with modifications to the protocol as follows: at day 0, the cells were plated at 5×10^5^ cell to 60 mm dish in serum-free differentiation (SF-D) medium (Nostro *et al*, 2008), and allowed to aggregate as embryoid bodies (EBs). After 48h, EBs were collected and dissociated into single cells by TrypLE Express Enzyme (Gibco). Single cells were plated at 1×10^5^ cell/well in 12-well plate in SF-D media containing recombinant human bone morphogenetic protein 4 (rhBMP4, 10 ng/ml, Qkine), Activin A (5 ng/ml, Qkine), and recombinant human vascular endothelial growth factor (rhVEGF, 5 ng/ml, Qkine). At day 5, EBs were washed and split 1:2 in SF-D media containing rhVEGF (5 ng/ml, Qkine) and stem cell factor (SCF, 50 ng/ml, Qkine). From day 5, EBs were cultured in 12-well plate coated with anti-adherence rinsing solution (STEM CELL TECHNOLOGIES).

### Flow cytometry

EBs were collected and dissociated into single cells in TrypLE Express Enzyme (Gibco). Cells were washed three times in FACS buffer (2% FCS in PBS), stained with antibodies diluted in FACS buffer for 30 min on ice. Samples were analysed using BD LSR Fortessa or CytoFLEX S (Beckman Coulter). Flow cytometry data was analysed using FlowJo software (BD Biosciences). Antibodies used for flow cytometry analysis were listed in Table S4.

### Immunofluorescence

Embryos were fixed in 4% paraformaldehyde (PFA) for 30 min at room temperature (RT), followed by washing in 0.5% polyvinylpyrrolidone in PBS. Embryos were permeabilized in 0.5% Triton X-100 in PBS for 15 min and blocked with 2% donkey serum, 2.5% BSA, and 0.1% Tween20 in PBS for 1 h at RT. For phosphorylated-STAT3 staining, permeabilization was performed in absolute methanol for 10 min at −20℃. Primary antibodies were diluted in blocking solution and incubated overnight at 4℃. After washing in 0.1% Tween 20 in PBS, embryos were incubated with Alexa Fluor-conjugated secondary antibodies (Thermo) for 1-3 h at RT. Nuclear staining was carried with Hoechst33342 (Thermo). Primary and secondary antibodies used are listed on Table S4.

For IF of ESCs and EpiSCs, cells were cultured on fibronectin coated ibidi-treated 8-well chamber slides (ibidi). Cells were fixed in 4% PFA for 15 min at RT, followed by washing in PBS. Cells were permeabilized in absolute methanol for 10 min at −20℃, then washed in PBS. Blocking reaction was carried out with 2% donkey serum, 2.5% BSA, and 0.1% Tween20 in PBS for 1 h at RT. Primary antibodies were diluted in blocking solution and incubated overnight at 4℃. After washing in 0.1% Tween 20 in PBS, cells were incubated with Alexa Fluor-conjugated secondary antibodies (Thermo) for 1 h at RT. Nuclear staining was carried with Hoechst33342 (Thermo).

### Imaging and image analysis for ESCs and EpiSCs

Images of ESCs and EpiSCs were acquired with LSM980 (Zeiss) confocal microscope and processed with ImageJ. Segmentation of nucleus and quantification of pSTAT3 levels were performed using CellProfiler 4.2.1 (McQuin *et al*, 2018). Mean intensity value of pSTAT3 in each nuclear section was normalized by mean intensity of Hoechst channel.

### Bulk RNA-sequencing

For low-input RNA-sequencing, embryos from E5.5 to E7.5 were dissected from *Stat3* heterozygous inter-crosses, and extraembryonic ectoderm and visceral endoderm removed manually using a flame-pulled Pasteur pipette of appropriate diameter. Extraembryonic ectoderm was used for genotyping. RNA was isolated from single epiblasts with Picopure RNA-isolation kit (Thermo). For library construction, SMARTerR Stranded Total RNA-seq kit v2-Pico InputMammalian (Takara Clontech) was used. For bulk RNA-sequencing for EpiSCs, total RNA was isolated using RNeasy Mini Kit (QIAGEN). Ribo-Zero rRNA Removal kit (Illumina) was used for rRNA removal and NEBNext Ultra II DNA Library Prep Kit for Illumina was used for the library construction. Reads were aligned to the mouse (GRCm38/mm10) reference genome using HISAT2(v2.2.1). StringTie (v2.1.4) was used for gene counts. R environment and Bioconductor were used for all RNA-seq analysis. Normalization and differential expression analysis were carried out with DESeq2 package (v1.30.1). Transcripts with absolute log2 fold change > 2 and the p-value adjusted (padj) < 0.05 considered as differentially expressed gene. Regularized log transform (rlog) normalized values were used for principal component analysis (PCA) and heatmap. GO-term enrichment analysis was performed with Metascape (Zhou *et al*, 2019). WT and *Stat3* null ESC bulk RNA-seq data was obtained from (Betto *et al*., 2021).

### Blastocyst injection of *Stat3* null ESCs

tdTomato-expressing Stat3 null ESCs were dissociated with Accutase and resuspended in 20 mM Hepes containing N2B27 medium. Around 10 cells were injected into each E3.5 CD1 blastocyst, incubated in N2B27 at 37℃ in 7% CO_2_ for ∼1 h to allow re-expansion, then transferred to pseudo pregnant female mice generated by mating to vasectomised males 2.5 days previously. Dissected embryos were imaged using a Leica stereo microscope. For sectioning, embryos were fixed in 4% PFA for overnight, which was replaced with 20% sucrose/PBS and incubated overnight at 4°C then embedded in OCT compound and sectioned at 8 μm thickness. Sections were imaged using Zeiss apotome microscope.

### Chimera-seq analysis

Sample preparation for Chimera-seq was performed as previously reported (Pijuan-Sala *et al*., 2019). Briefly, chimeric embryos were collected at E7.5, E8.5, and E9.5 and pooled in 1.5 ml tubes (∼10 embryos for E7.5, ∼ 4 embryos for E8.5, 1 embryo for E9.5). Embryos were dissociated into single cells by TrypLE™ Express Enzyme, then tdTomato+ and tdTomato-cells were sorted using a BD Influx sorter for subsequent 10x scRNA-seq library preparation and sequencing on an Illumina HiSeq 4000 platform.

### scRNA-seq pre-processing

Raw sequencing files were mapped to the mm10 reference genome and counted with the GRCm38.p5 annotation including the Tomato-Td gene using CellRanger count (v6.0.1) with chemistry=SC3Pv3. All downstream processing and analysis were performed in R (v4.2.2). High quality cells were retained using the following thresholds: log10(number of reads) > 4 & < 5, number of genes > 2e3 & < 1e4, percentage mitochondrial RNA < 5%, percentage ribosomal RNA < 30%. Count normalization was performed by first calculating size factors using the computeSumFactors function from scran (Lun *et al*, 2016), where cells were pre-clustered using scran’s quickCluster with method=igraph, minimum and maximum sizes of 100 and 3000 cells per cluster, respectively, followed by logNormCounts from scuttle (McCarthy *et al*, 2017). Doublet calling was performed with the scds (Bais & Kostka, 2020) package using the hybrid approach and doublet score > 1 as threshold. Accidental sorting of Stat3 KO cells into the WT samples was corrected by annotating whether a cell from a WT sample contained any read mapping to the Tomato-Td gene, and those cells were excluded from downstream differential abundance and differential expression testing.

### Label transfer from extended transcriptional atlas of mouse gastrulation and early organogenesis

Label transfer was performed for cell-type assignment by mapping to the E7.5 – E9.5 stages of the extended gastrulation atlas (Imaz-Rosshandler *et al*., 2024). First, the reference atlas was subset to a maximum of 15k cells per embryonic stage and all cells for the ‘mixed-gastrulation’ stage. Mapping was then performed for every sample separately using the batchelor package (Haghverdi *et al*, 2018), each sample is referred to as the ‘query’. Joint normalisation was performed by first running MultiBatchNorm for the reference atlas with samples as batch, followed by cosineNorm over the logcounts of both the reference atlas and the query. For downstream analysis, non-informative and noisy genes were removed, these include the genes which names start with Rik, Mt, Rps, Rpl, or Gm, end with Rik, haemoglobin genes, imprinted genes, Xist and Tsix, Tomato-Td, and Y-chromosomal genes. This was followed by selecting the top 2,500 highly variable genes in the reference atlas using modelGeneVar (Scran) with samples as block, which are used for downstream analysis. Dimensionality reduction was performed using MultiBatchPCA for the concatenated query and reference atlas objects, with the ‘atlas’ and ‘query’ as batch, and d=40. Batch correction was performed in multiple rounds using ReducedMNN, where first the atlas was corrected within each embryonic stage with samples ordered from largest to smallest, followed by atlas correction between embryonic stages with embryonic stages ordered from latest to earliest. This was followed by batch correction between the reference atlas and the query. Nearest neighbours (NN) of every query cell in the reference atlas were identified using queryKNN from the BiocNeighbors (v1.8.2) package with k=25. Cell type label transfer was then performed by taking the mode of the cell types of the NN, ties were broken by taking the cell-type of the reference atlas cell closest to the query.

### Dimensionality Reduction

Dimensionality reduction was performed by first detecting the top variable genes in the same way as described in the ‘Label transfer from extended transcriptional atlas of mouse gastrulation and early organogenesis’ section. Next, the number of genes and reads were regressed out as covariates in the log-normalised count matrix. Next, PCs were calculated using the prcomp_irlba function from the irlba (v2.3.3) package, with n=40. Batch correction was performed as described for the reference atlas above, correcting both within and between embryonic stages. Uniform Manifold Approximation and Projection (UMAP) was performed using Scater’s runUMAP function on batch-corrected PCs, with n_neighbors=30 and min_dist=0.3.

### Differential abundance testing

Differential abundance testing was performed for each embryonic stage separately using the MiloR package (Dann *et al*, 2022). First, embryonic stage specific dimensionality reduction and batch correction was performed as described above, except for the between embryonic stage batch correction step. Next, Milo was performed on embryonic stage specific batch corrected PCs by first running buildGraph with k=30 followed by makeNhoods with k=30 and prop=0.05. Cells per neighbourhood were counted using countCells and differential abundance testing was performed using testNhoods with design = ∼ embryo pool + genotype, where each embryo pool contains one sample for both the Stat3 KO and the WT condition that were collected as chimeric embryos and separated by the presence or absence of tdTomato expression by fluorescence-activated cell sorting.

### Cell type differential gene expression testing

Differential gene expression testing was performed on the subset of genes after filtering out non-informative and noisy genes as described in the ‘Label transfer from extended transcriptional atlas of mouse gastrulation and early organogenesis’ section. Next, cells were pseudobulked by their label transferred cell type, original sample, and genotype, where only those conditions with more than 15 cells were retained. Differential genes were then detected using Scran’s pseudoBulkDGE (using edgeR v3.32.1) function, with the label transferred cell types, the design the same as for the testNhoods function in ‘Differential abundance testing’, and the genotype as the condition. The significance thresholds were set at a FDR<0.05 and log2 fold change > 1.

### Erythroid trajectory

Cells labelled as ‘Hematoendothelial progenitors’, ‘Blood progenitors’, or ‘erythroid differentiation trajectory’ were used for inference of the erythroid trajectory. Gene filtering and top 1,000 highly variable gene selection was performed as described in the ‘Label transfer from extended transcriptional atlas of mouse gastrulation and early organogenesis’ section. PCs were calculated using Scater’s runPCA with ncomponents=5 and PCs were used to generate the diffusionmap using the DiffusionMap function from the destiny package (Angerer *et al*, 2016). Pseudotime was estimated using slingshot (Street *et al*, 2018) with PCs as input. For differential expression testing along pseudotime, only cells with a pseudotime value below 50 were used to have comparable densities for both genotypes. Genes were filtered as described above, with the additional filter of genes having to be detected in at least 10% of cells in either of the genotypes. Differential expression testing was performed using the TradeSeq package (Van den Berge *et al*, 2020). First, a negative binomial generalized additive model was fitted for every gene using fitGAM with condition=‘genotype’, U=‘embryo pool’, pseudotime=‘slingshot pseudotime’, nknots=6, and cellWeights set to ‘1’ for every cell as all cells belong to the same trajectory. Next, differential genes were detected using conditionTest with l2fc=1. Multiple-comparison correction was done using p.adjust with method=‘fdr’ and genes with corrected p-values of <0.05 were labelled as significant.

## Data Availability

Bulk RNA-seq data and chimera-seq data are deposited in Gene Expression Omnibus (accession no GSE260590).

## Acknowledgements

We wish to thank Kenneth Jones for lab management and help with genotyping; Thomas Burdon and Lawrence Bates for helpful discussion on the project; Shizuo Akira for providing *Stat3+/-* mice; Peter Humphreys and Darran Clement (Cambridge Stem Cell Institute), Ann Wheeler and Laura Murphy (Human Genetics Unit, University of Edinburgh) for advanced imaging and analysis; Michael Rennie and Elizabeth Freyer for flow cytometry analysis; Maike Paramor and Vicky Murray for sequencing (Cambridge Stem Cell Institute); UBS, Cambridge for animal husbandry; So Shimamoto and Shota Nakanoh for helpful discussion. This work was supported by the University of Cambridge, BBSRC project grant RG74277 and funded in part by Wellcome Trust (Grant number: 203151/Z/16/Z). T.A. was supported by JSPS Overseas Research Fellowship and UEHARA Memorial Foundation Fellowship.

The authors declare no competing interests

---

**Fig. S1.**
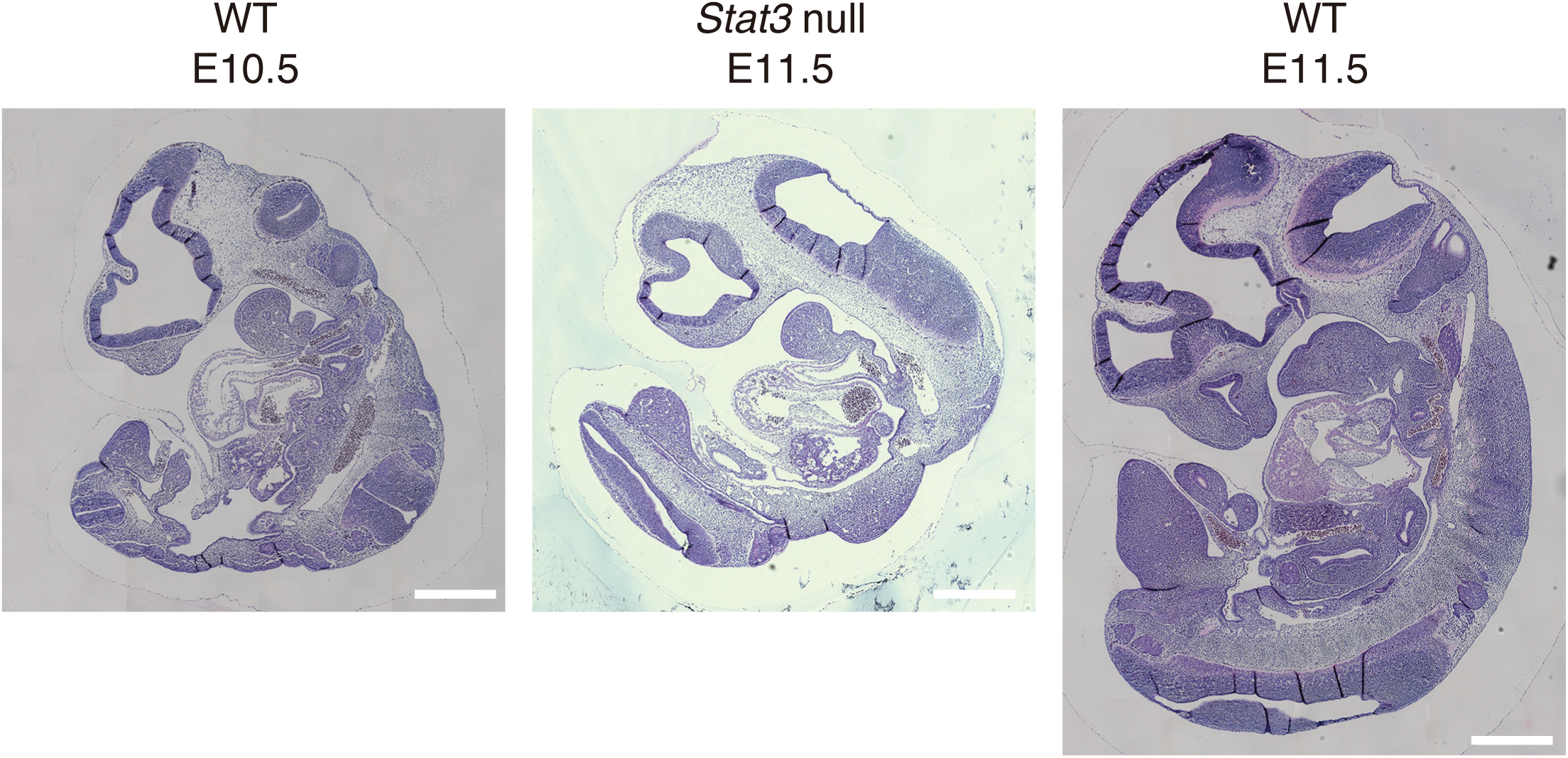
Histological sections for WT and *Stat3* null embryos. Hematoxylin and eosin staining for E10.5/11.5 WT and E11.5 Stat3 null embryos. Scale bar = 500 µm.

**Fig. S2.**
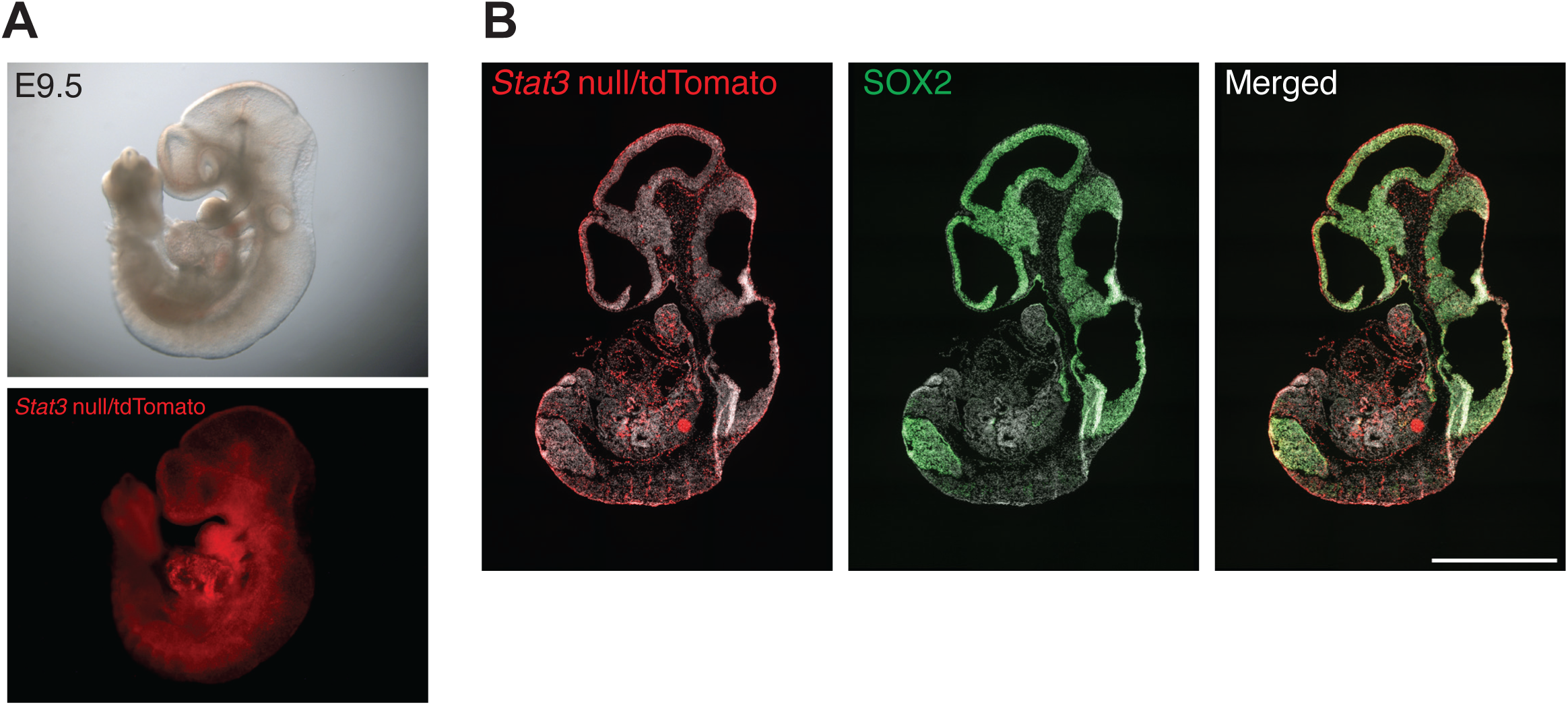
Contribution of *Stat3* null cells in chimeric embryos. (A) Bright field (top) and tdTomato fluorescent (bottom) images of chimeric embryos. tdTomato-expressing *Stat3* null ESCs were injected host blastocyast and chimeric embryos were collected at E9.5. (B) Immunofluorescence for SOX2 in chimeric embryos. Note tdTomato-positive *Stat3* null cells contributed to multiple tissues in the embryo, including SOX2-positive neural lineages in the brain.

**Fig. S3.**
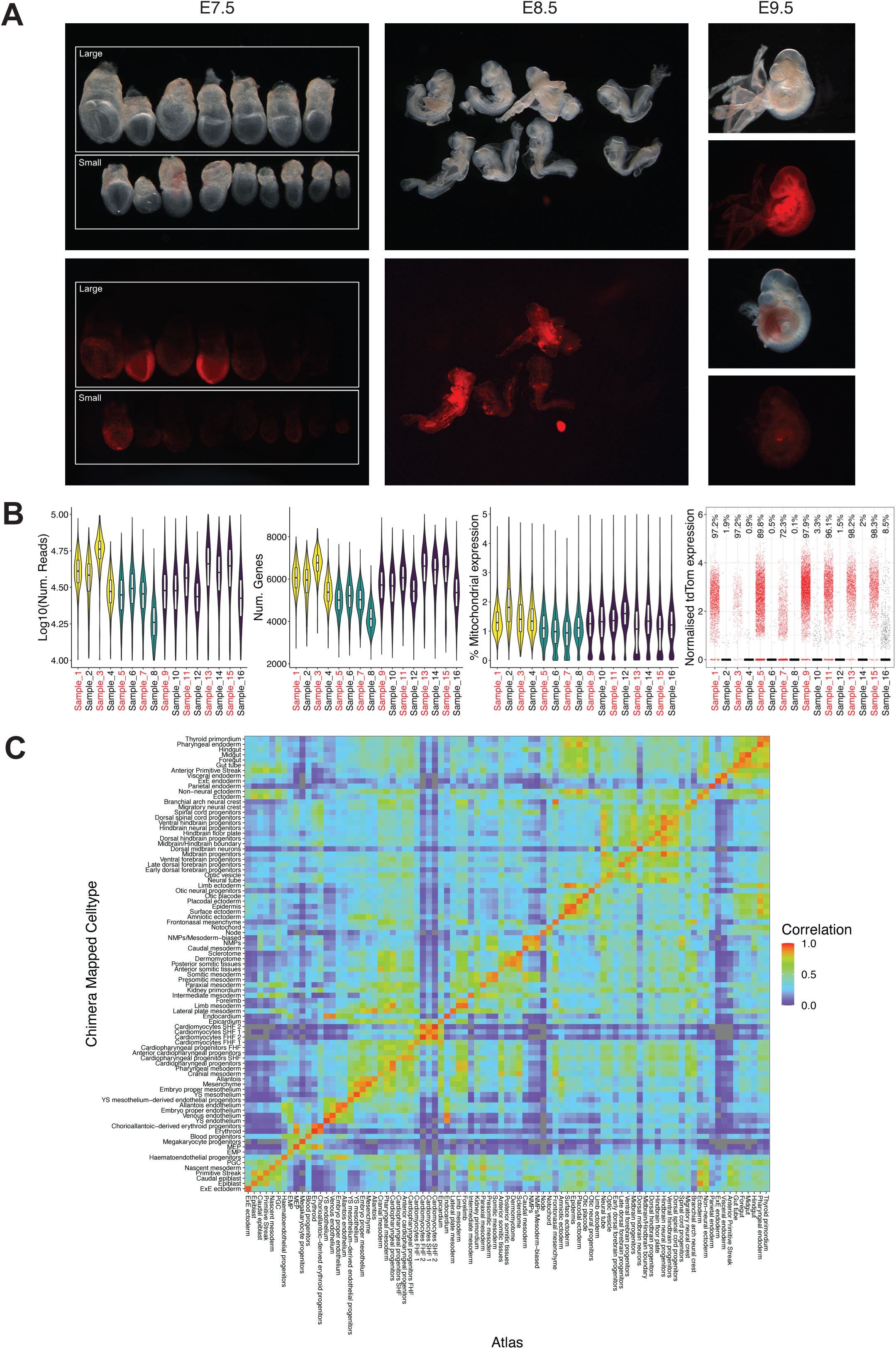
Chimera embryos used for single-cell Chimera-seq and quality control of single-cell chimera-seq. (A) Bright field and tdTomato fluorescent images for E7.5, E8.5, and E9.5 chimeric embryos used for single-cell Chimera-seq. For E7.5, embryos were classified by their size (large and small). 7 large and 9 small E7.5 embryos were pooled and processed for scRNA-seq. For E8.5, 3 or 4 embryos were pooled and processed for scRNA-seq. For E9.5, two embryos were processed individually for scRNA-seq. (B) Quality of chimaera-seq cells per sample after filtering, showing number of reads, number of genes, percentage of mitochondrial expression, and normalised tdTomato expression including percentage of cells that are non-zero. Stat3 KO and WT samples are in red and black, respectively. E7.5 – yellow, E8.5 – green and E9.5 – purple. (C) Heatmap showing Pearson correlation of expression of cell type marker genes between cell types of the extended transcriptional atlas of mouse gastrulation and early organogenesis and label transferred cell types of the Stat3 KO chimera.

**Fig. S4.**
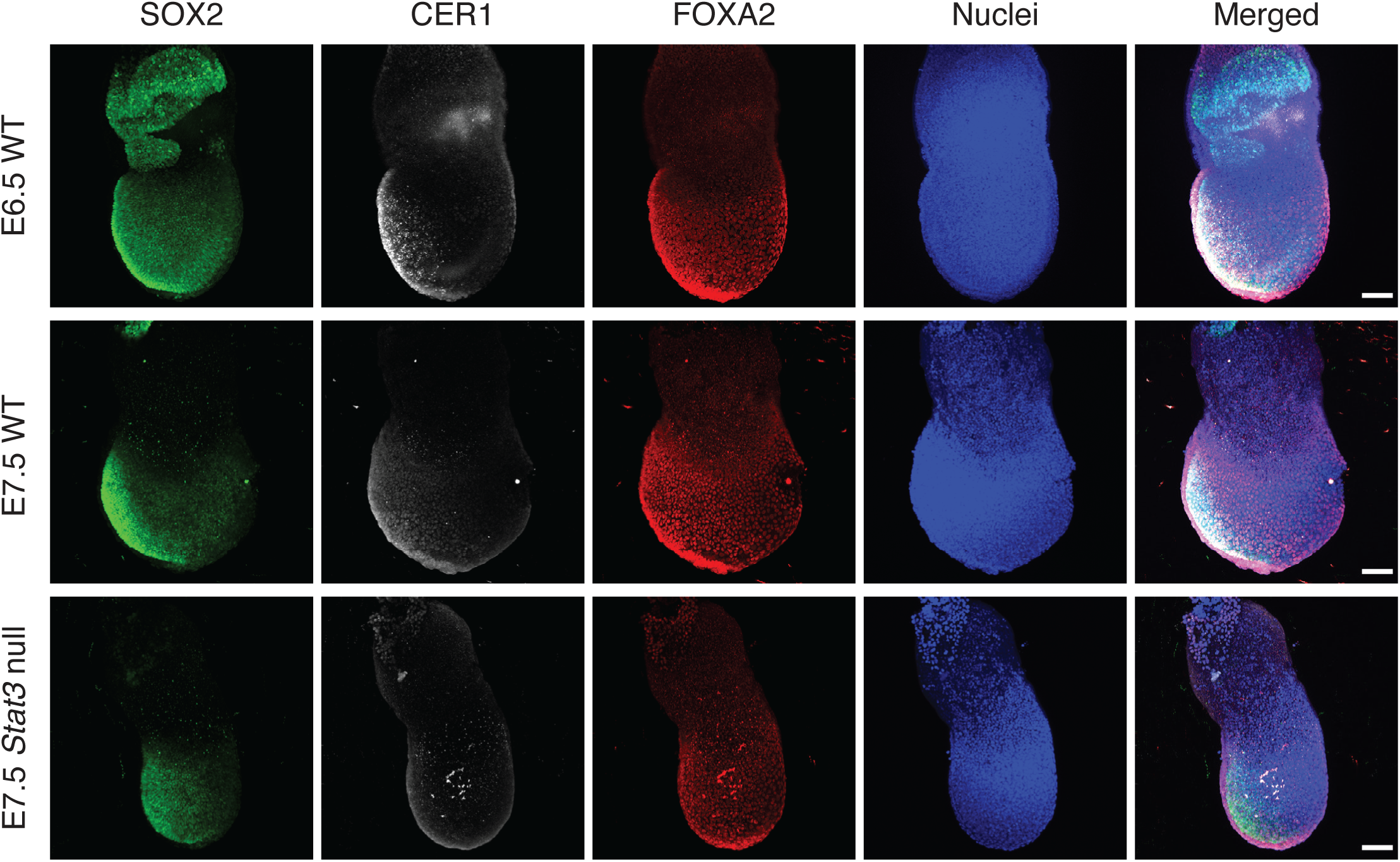
Extra-embryonic endoderm development in *Stat3* null embryos. Immunofluorescence for SOX2, CER1, and FOXA2 in E6.5/7.5 WT and E7.5 *Stat3* null embryos. E6.5 WT (n = 5), E7.5 WT (n = 6), and E7.5 *Stat3* null (n = 3) were examined. Scale bar = 50 µm.

**Fig. S5.**
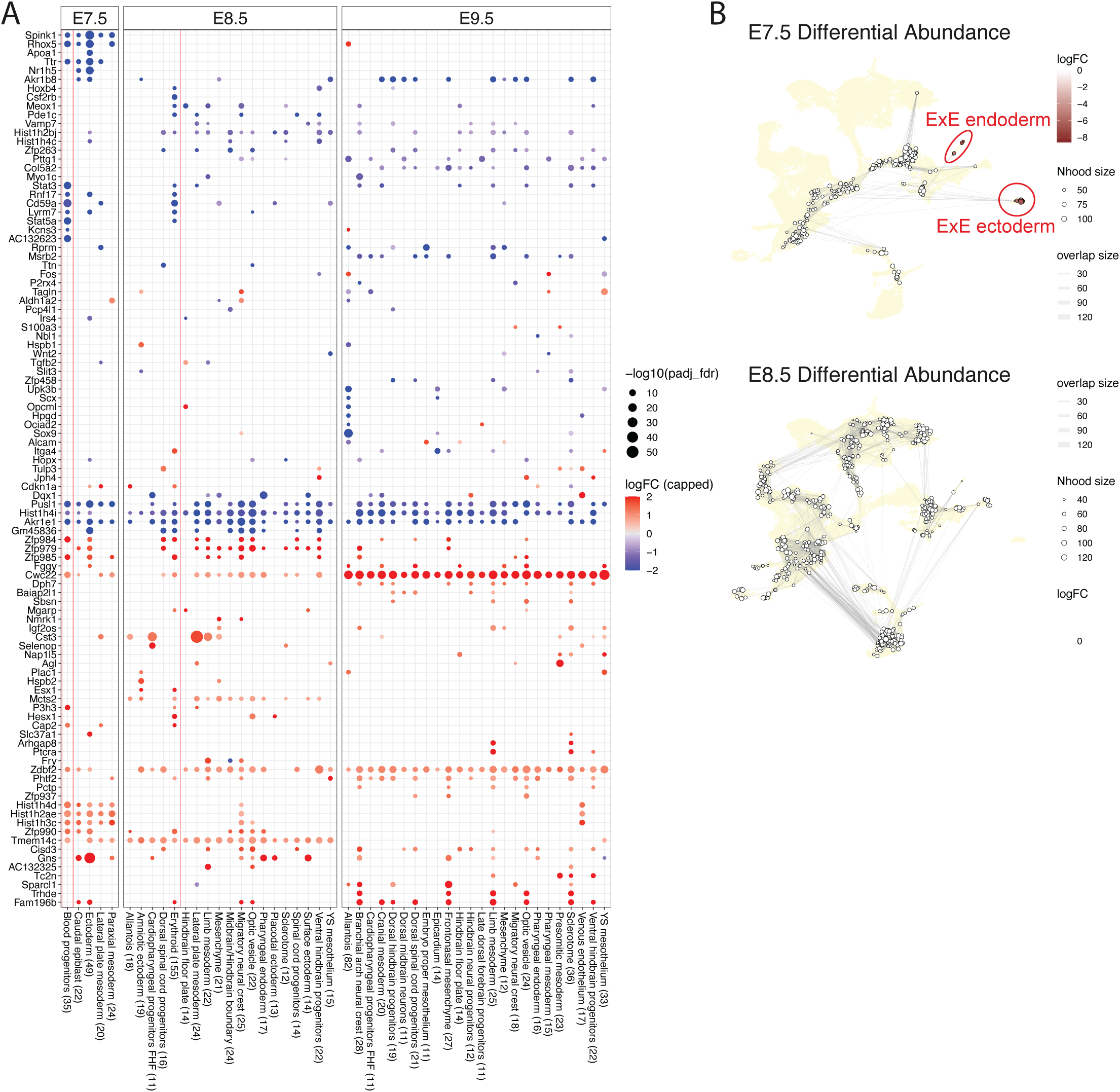
Differential expression and abundance of *Stat3* null and WT cells in E7.5, E8.5, and E9.5 chimeras. (A) Differential expression between *Stat3* null and WT cells for label transferred cell types at each embryonic stage. Only cell types with more than 10 differential genes and genes differentially expressed in more than 1 cell types are shown. Positive values represent genes that are more highly expressed in *Stat3* null. (B) Differential abundance testing between *Stat3* null and WT cells from E7.5 and E8.5 chimera-seq data. Negative logFC indicates a loss of *Stat3* null cells in a specific cellular neighbourhood.

**Fig. S6.**
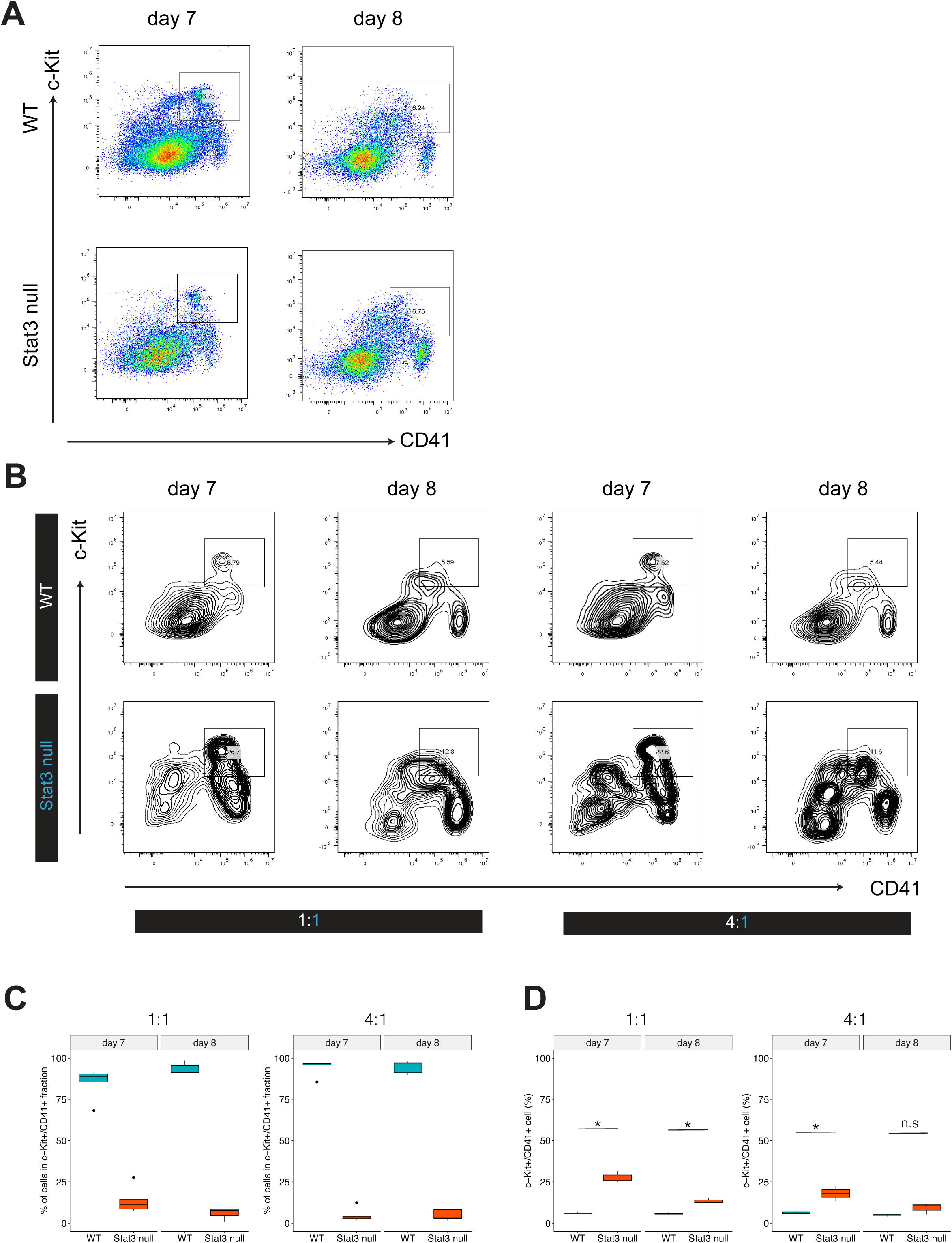
Hematopoietic progenitor differentiation from *Stat3* null ESC. (A) Representative flow cytometry analysis of CD41/c-Kit expression in day 7 and 8 WT and *Stat3* null EBs. CD41^+^/c-Kit^+^ cells are gated in the plot. n = 8 (WT) and n = 8 (*Stat3* null) independent differentiation. (B) Representative flow cytometry analysis of CD41/c-Kit expression at day 7 and day 8 in WT and *Stat3* null cells in chimeric EBs, mixing WT and *Stat3* null ESCs at 1:1 and 4:1. n = 3 independent differentiation. (C) Percentages of WT and *Stat3* null cells in the c-Kit^+^/CD41^+^ fraction at day 7 and day 8. (D) Percentages of c-Kit^+^/CD41^+^ fraction at day 7 and day 8 in WT and *Stat3* null cells in chimeric EBs, gated in (B). n.s, not significant. * *p* < 0.05.

**Fig. S7.**
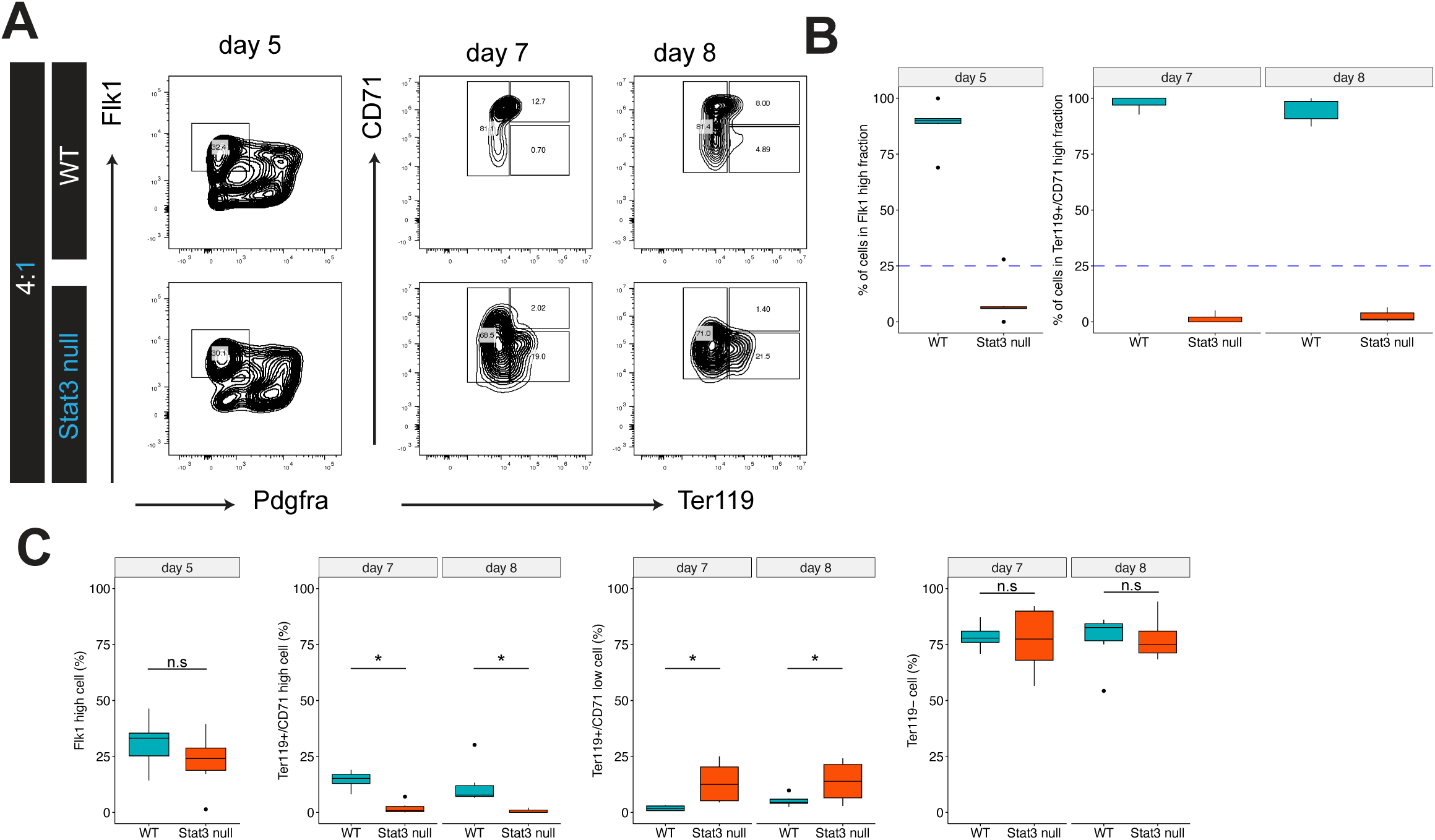
Primitive erythroid differentiation in chimeric EBs. (A) Representative flow cytometry analysis of Flk1/Pdgfra expression in day 5 and Ter119/CD71 expression in day 7 and day 8 in WT and *Stat3* null cells in chimeric EBs, mixing WT and *Stat3* null ESCs at 4:1. (B) Percentages of WT and *Stat3* null cells in the Flk1^hi^ fraction at day 5, and in the Ter119^+^/CD71^hi^ fraction at day7 and day 8. (I) Percentages of Flk1^hi^ cells at day 5, and Ter119^+^/CD71^hi^, Ter119^+^/CD71^low^, and Ter119^−^ cells at day 7 and day 8 in WT and *Stat3* null cells in chimeric EBs, gated in (A). n.s, not significant. * *p* < 0.05.

**Table S1.** Numbers of *Stat3* WT, Het, and null embryos from *Stat3* heterozygous intercrosses.

**Table S2.** EpiSCs derivation from epiblast of *Stat3* heterozygous intercrosses.

**Table S3.** List of primers for genotyping.

**Table S4.** List of primary and secondary antibodies.

**Supplementary Table 1.** Genotypes of embryos from Stat3+/− intercross.

|  | **+/+ or +/-** | **-/-** | **Empty decidua** |
| --- | --- | --- | --- |
| **E3.5** | 50 | 16 |  |
| **E4.5** | 76 | 21 |  |
| **E5.5** | 22 | 6 |  |
| **E6.5** | 63 | 18 | 2 |
| **E7.5** | 45 | 10 | 4 |
| **E8.5** | 35 | 5 | 4 |
| **E9.5** | 42 | 2 | 5 |
| **E10.5** | 18 | 1 | 3 |
| **E11.5** | 29 | 4 | 3 |

**Supplementary Table 2.** EpiSC deriva6on from epiblast of Stat3+/- intercross.

| Embryo stage | Lines established |  | Number of<br>epiblast plated | Efficiency |
| --- | --- | --- | --- | --- |
|  | +/+ or +/- | -/- |  |  |
| E6.5 | 14 | 3 | 17 | 100% |
| E7.5 | 10 | 3 | 13 | 100% |

**Supplementary Table 3.**
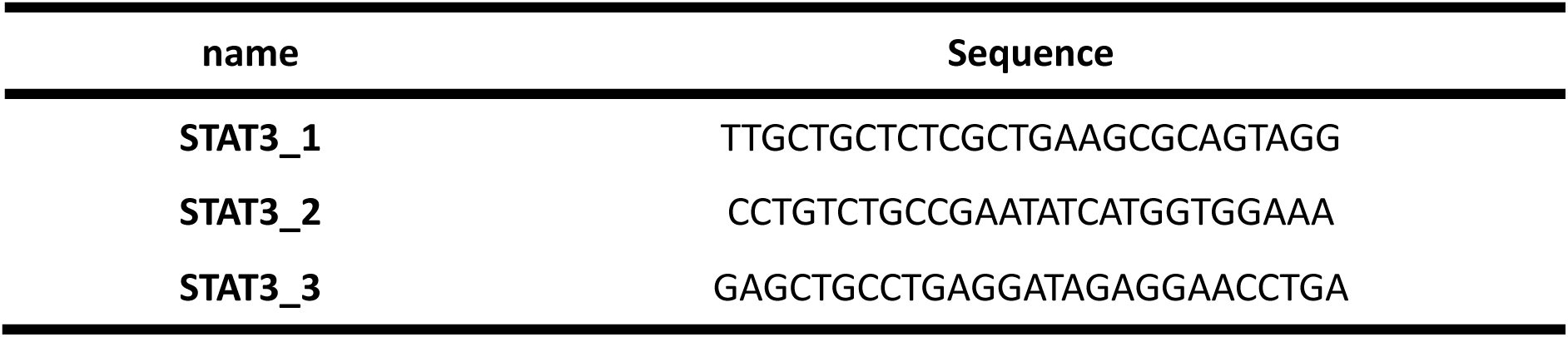
List of genotyping primers.

**Supplementary Table 4.** List of an6bodies.

| Antibodies | Source | Cat# |
| --- | --- | --- |
| Mouse monoclonal anti-Oct3/4 (C-10) | Santa Cruz | Cat#SC-5279;<br>RRID:AB_628051 |
| Goat polyclonal anti-Brachyury | R&D systems | Cat#AF2085;<br>RRID:AB_2200235 |
| Rat monoclonal anti-Nanog | eBioscience | Cat#14-5761-80;<br>RRID:AB_763613 |
| Rat monoclonal anti-OCT3/4 | eBioscience | Cat#14-5841-82;<br>RRID:AB_914301 |
| Rat monoclonal anti-SOX2 | eBioscience | Cat# 14-9811-82,<br>RRID:AB_11219471 |
| Rabbit anti-phospho-(Tyr705) STAT3 | Cell Signaling Technology | Cat#9145;<br>RRID:AB_2491009 |
| Rabbit anti-phospho-(Ser727) STAT3 | Cell Signaling Technology | Cat#34911;<br>RRID:AB_2737598 |
| Goat anti-GATA6 | R&D systems | Cat#AF1700;<br>RRID:AB_2108901 |
| Goat anti-Brachyury | R&D systems | Cat#AF2085;<br>RRID:AB_2200235 |
| Goat anti-CER1 | R&D systems | Cat#AF1075;<br>RRID:AB_2077228 |
| Rabbit anti-FOXA2 | abcam | Cat# ab108422;<br>RRID:AB_11157157 |
| BV786 conjugated anti-Flk1 | BD Biosciences | Cat# 740938<br>RRID:AB_2740568 |
| PE conjugated anti-Pdgfra | eBioscience | Cat# 12-1401-81<br>RRID:AB_657615 |
| PE conjugated anti-CD71 | eBioscience | Cat# 12-0711-82<br>RRID:AB_465740 |
| FITC conjugated anti-Ter119 | eBioscience | Cat# 11-5921-82<br>RRID:AB_465311 |
| PE-Cy7 conjugated anti-CD41 | eBioscience | Cat# 25-0411-82<br>RRID:AB_1234970 |
| APC conjugated anti-c-Kit | eBioscience | Cat# 17-1171-82<br>RRID:AB_469430 |
| Donkey anti-rat Alexa488 | Thermo | Cat# A-21208,<br>RRID:AB_2535794 |
| Donkey anti-mouse Alexa488 | Thermo | Cat# A-21202,<br>RRID:AB_141607 |
| Donkey anti-rabbit Alexa555 | Thermo | Cat# A-31572,<br>RRID:AB_162543 |
| Donkey anti-goat Alexa647 | Thermo | Cat# A-21447,<br>RRID:AB_2535864 |
| Donkey anti- Mouse Alexa647 | Thermo | Cat# A-31571,<br>RRID:AB_162542 |

